# Consistent drought regulation in grapevine is driven by directional transcription factor activity

**DOI:** 10.1101/2025.11.14.688560

**Authors:** G Vásquez-Marambio, TC Moyano, D Navarro-Payá, A Sequeida, F Gaínza-Cortés, T Matus, A Orellana, JM. Álvarez

## Abstract

Climate change is intensifying environmental stresses such as drought, threatening vineyard productivity and sustainability worldwide. *Vitis vinifera* cultivars, responsible for most wine and table grape production, are particularly sensitive to water deficit, whereas many rootstocks derive from different *Vitis* species or interspecific hybrids with higher stress tolerance. A key step toward mitigating the effects of severe drought is the identification of regulatory genes controlling drought responses, enabling the design of gene expression–based strategies or the generation of resilient cultivars through new breeding technologies. In this study, we performed a meta-transcriptomic analysis to identify genes consistently differentially expressed under drought in cultivated *V. vinifera* and two hybrid rootstocks (M4 and 101-14). Using more than twenty drought-control comparisons, we identified a core set of 4,617 drought-responsive genes that were consistently mis regulated across multiple experimental conditions. This core gene set was used to construct gene regulatory networks integrating genome-wide transcription factor (TF) binding motif analysis with random forest-based regulatory network generation employing machine learning techniques. We identified key TFs, including the Abscisic-Acid-(ABA) Responsive Element Binding Factor 2 (ABF2), MYB30A and an uncharacterized HMGbox domain protein, as central regulators within the network. Several top-ranking TFs, displaying up-or down-regulation under drought conditions, were primarily identified as positive regulators of their target genes, while lower-hierarchy TFs exhibited inverse expression relationships with their predicted targets. The network exhibited a hierarchical organization architecture among several TFs whose homologues in other species are linked to ABA signaling, with several TF families represented, each potentially operating at distinct regulatory tiers. Some TFs appear to act as central hubs orchestrating broad transcriptional programs, whereas others likely control more specialized branches of the drought response. Overall, these findings offer novel insights into the transcriptional control of drought tolerance in grapevine and provide key candidate regulators for breeding and biotechnological strategies aimed at improving stress resilience.

## 1. Introduction

Grapevine (*V. vinifera*) is one of the most economically important fruit crops worldwide, cultivated across approximately 7.2 million hectares (OIV, 2023). However, grapevine productivity and fruit quality are highly sensitive to environmental stresses, with drought representing one of the most detrimental factors limiting agricultural production (FAO, 2023). Climate projections indicate that drought events will become more frequent and severe in many regions across the globe (IPCC, 2023), posing a growing challenge for viticulture.

Drought stress leads to significant changes in plant systems, including physiological, biochemical, and molecular alterations that contribute to decreased growth, productivity, and quality (Cattivelli et al., 2008). In response to drought, plants modify gene expression patterns to cope with drought through Gene Regulatory Networks (GRNs) (Singh & Laxmi, 2015). GRNs are dynamic and hierarchical systems composed of regulatory interactions, where TFs act as master regulators (Song et al., 2016). By binding to specific cis-regulatory elements in the promoters of their target genes, TFs orchestrate gene expression programs that enable plants to adapt to water deficit conditions (Yamaguchi-Shinozaki & Shinozaki, 2006).

GRN inference has emerged as a strategy to unravel these complex regulatory circuits, particularly in crops, where experimentally validated TF–target interactions are limited. By combining transcriptomic data with computational approaches, GRN inference allows the prediction of regulatory relationships and the identification of key TFs involved in drought responses (Kulkarni et al., 2019). This systems-level understanding is critical for guiding genetic improvement strategies aimed at enhancing drought tolerance.

Considerable advances have been made in identifying drought-responsive genes in crops, facilitated by transcriptomic approaches such as RNA-seq (Joshi et al., 2016). These genes encode proteins involved in metabolic processes, including detoxification, osmolyte biosynthesis, and ion transport, which help the plant cope with water scarcity (Joshi et al., 2015). Among these drought-responsive genes, a particularly important group consists of TFs, including members of the ABF, AP2/ERF, NAC, bZIP, MYC, and MYB families (Wani et al., 2013). These TFs regulate stress-responsive genes by binding to promoter regions and modulating transcription in response to drought (Franco-Zorrilla et al., 2014). In addition, signaling molecules such as MAPKs, CDPKs, and protein phosphatases coordinate with TFs to fine-tune stress signaling and gene expression (Joshi et al., 2016). However, most of our current understanding of drought-responsive GRNs is derived from studies in model species such as *Arabidopsis thaliana*, leaving significant knowledge gaps regarding the regulatory networks in woody crops like *V. vinifera*. While several TFs have been characterized in grapevine stress responses, the overall transcriptional hierarchy within the GRN, remains to be characterized. This represents a major obstacle to developing effective strategies for improving drought tolerance in grapevine.

Gene co-expression networks (GCNs), a type of GRN analysis, have been used in *V. vinifera* to investigate a wide range of biological processes. These include responses to biotic stresses, such as pathogen resistance (T. Li et al., 2024), and fruit quality formation (Fu et al., 2023). GCNs have also been applied to the study of abiotic stresses, including heat and drought (Tan et al., 2023; Wu et al., 2023) and specialized metabolic processes (Orduña et al., 2023). Despite the valuable insights provided by GCNs into grapevine biology, their inherent limitations restrict their capacity to fully resolve transcriptional regulatory mechanisms. GCNs are based on correlation, and as such, they do not provide evidence of causality nor directionality between genes. This limits the ability to distinguish whether a transcription factor is directly regulating a target gene or simply co-expressed with it (B. Zhang & Horvath, 2005). Another disadvantage is that correlation networks do not necessarily distinguish between direct and indirect (i.e., downstream) TF-target interactions, potentially leading to many false-positive predicted interactions (Xiao et al., 2022). To address these limitations, integrative GRN inference approaches have been developed. These methods combine gene expression data with sequence-based information, such as cis-regulatory motif analysis, to infer more accurate and biologically meaningful TF-target interactions (Gupta et al., 2021).

In addition, integrative genomics approaches enable the detection of consistently regulated genes across heterogeneous experimental datasets, encompassing different conditions, tissues, and developmental stages. These “condition-proof” genes are more likely to represent robust targets for improving drought tolerance, as they function reliably across diverse scenarios (Buti et al., 2019; Muthuramalingam et al., 2017; Shahriari et al., 2022). This strategy enhances the likelihood of identifying key regulatory components that are effective under variable environmental conditions, which is essential for breeding programs and genetic engineering aimed at enhancing crop resilience.

In this study, we applied an integrative genomics approach to infer a comprehensive GRN of the drought response in *V. vinifera*. By analyzing transcriptomic datasets from multiple experimental conditions and integrating expression patterns with cis-regulatory motif information, we identified key master regulators orchestrating the drought response. Our analysis reveals that TFs such as ABF2, MYB30A, bHLH044, HMGbox(u), and PIF4 show high regulatory potential. These TFs function as central hubs within the inferred GRN and exhibit high regulatory influence over drought-responsive genes involved in stress adaptation pathways, including osmotic adjustment, reactive oxygen species detoxification, and hormone signaling. Importantly, we also examined the directionality of regulatory interactions, revealing that upregulated TFs tend to activate upregulated targets, while TFs whose expression is downregulated are often linked to downregulated targets. In addition, TFs themselves tend to interact with other TFs showing similar expression patterns. The identification of these master regulators and their predicted target genes provides novel insights into the transcriptional control of drought tolerance in grapevine. This coordinated directionality suggests that drought responses in grapevine are orchestrated through a structured and coherent regulatory hierarchy, where transcriptional programs are either globally activated or repressed. By offering a systems-level perspective of GRN architecture, our findings lay a solid foundation for future functional validation studies and the development of drought-tolerant *V. vinifera* varieties. Such advances are critical for addressing the growing challenges posed by climate change in viticulture and ensuring the sustainability of grape production.

## 2. Materials and Methods

### 2.1. Selection and analysis of transcriptomic libraries

Using the search criteria “(Drought OR water-deprivation OR water deprivation) AND (txid29760[orgn] OR *Vitis vinifera* OR grape)”, we identified bioprojects of *V. vinifera* containing a total of 1180 transcriptomic libraries. Metagenomics libraries, transgenic lines, and libraries lacking available Control/Treatment data were excluded from the analysis. Ultimately, 550 libraries were selected and downloaded from the NCBI Sequence Read Archive (SRA) (https://www.ncbi.nlm.nih.gov/sra). Runs were trimmed and filtered with TrimGalore (Krueger et al., 2023, https://github.com/FelixKrueger/TrimGalore), pseudo-aligned against the PN40024 reference genome assembly (12X.v2) for *V. vinifera* (Canaguier et al., 2017) and quantified at the gene level with vCOST.v3 annotation, using Kallisto software (Bray et al., 2016). Differential expression analysis was performed using DESeq2 (Love et al., 2014) and genes with an adjusted p-value < 0.05 were considered differentially expressed genes. Gene symbols were extracted from the Grape Gene Reference Catalogue version 3 (Navarro-Payá et al., 2022).

### 2.2. Identification of Consistently Regulated Genes

To identify consistently regulated genes, we first collapsed all comparisons within each bioproject, including time-course experiments, into a single list of DEGs per bioproject. This step aimed to reduce redundancy and avoid overrepresentation from individual studies. Genes were then grouped according to the number of distinct bioprojects in which they were differentially expressed (i.e., present in at least three, four, or five bioprojects). For each group, we applied a hypergeometric test to evaluate enrichment for the Gene Ontology term ‘GO:0009414, Response to water deprivation’, using as background the total number of DEGs. The group of genes present in at least three bioprojects yielded the most significant p-value, suggesting that this threshold best captures genes consistently regulated across independent experiments. Detailed p-values for each group are provided in Supplementary Table 1 (ST1: Enrichment analysis of Water Deprivation-related DEGs across grouped bioprojects).

### 2.3 Gene Ontology Analysis

Gene Ontology enrichment analysis was conducted for consistently regulated genes using TopGO package (Alexa & Rahnenführer, 2023) using functional annotation files obtained from EggNOG mapper (Huerta-Cepas et al., 2017) (retrieved on 02/12/24).

### 2.4 Construction of gene regulatory networks

Transcription factor information for *V. vinifera* was obtained from the CISBP database (Catalogue of Inferred Sequences Binding Preferences, http://cisbp2.ccbr.utoronto.ca/) (Weirauch et al., 2014). Position Weight Matrix files were converted to MEME format using the Matrix2Meme tool from MEMEsuite (Bailey et al., 2015). Promoter regions were extracted with Bedtools (Quinlan & Hall, 2010), considering 2 kb upstream from the transcription start site (TSS). Gene regulatory networks were constructed using a genome-wide, motif-based analysis of promoter regions with Find Individual Motif Occurrences (FIMO) software (Grant et al., 2011). The GENIE3 machine learning algorithm was applied for integrating co-expression when inferring regulatory networks, filtering by the top 10% of interactions (Huynh-Thu et al., 2010). The total count matrix, normalized and processed with Combat Seq to remove batch effects, was used as input, along with a vector specifying the regulatory genes (Y. Zhang et al., 2020). The FIMO and GENIE3 networks were then merged to form a general GRN (i.e., merged network). A final drought-responsive network was constructed by selecting all interactions involving the set of consistent drought-responsive genes (drought responsive GRN).

### 2.5. Ranking TFs in the drought response

For upregulated and downregulated TFs, an average score was calculated by considering four criteria:

The magnitude of the response, assessed by comparing the absolute log2 fold changes between drought and control conditions, prioritizing TFs that exhibited the largest changes in expression levels. 2. Consistency across datasets, evaluated by determining the presence and direction of regulation in each drought/control contrast. 3. Regulatory influence, expressed as the proportion of drought-responsive target genes by each TF. This was determined through the outdegree of each TF in the gene regulatory network, which is a proxy of the number of target genes each TF regulates. For example, if the drought-responsive network contains 1,500 genes and a TF is predicted to regulate 120 of them, its regulatory influence would correspond to 8% of the network (120/1500). 4. Finally, statistical validation for specificity, calculated using a hypergeometric test, considering genome size, the size of the drought-responsive network, and each TF’s outdegree within both the merged and the drought-responsive GRN, with a false discovery rate (FDR) threshold of 0.01. The average score between all these criteria allowed for hierarchical ranking of the TFs based on their overall regulatory potential within the drought-responsive gene network. Transcription factors identified in the upregulated or downregulated rankings that exhibited at least three opposing regulations across drought/control contrasts were reclassified into the variable-response category. For the variable-response TF ranking, three key criteria were used: consistency across datasets, regulatory influence, and statistical validation. The average score for this category was calculated solely on these three criteria, as variable-response TFs displayed dynamic, condition-dependent expression patterns that warranted a distinct approach to their prioritization.

### 2.6 Transcription factor target gene enrichment analysis

To evaluate the functional relevance of transcription factor target genes in the drought responsive GRN, we performed a hypergeometric enrichment analysis using Gene Ontology (GO) categories associated with stress responses. Specifically, we assessed the enrichment of TF targets in GO:0009414 (Response to Water Deprivation), GO:0009737 (Response to Abscisic Acid), and GO:0009652 (Response to Salt Stress). Gene lists for each GO category were compiled from functional annotations. For each TF, we tested whether its target genes were significantly overrepresented in each GO category by comparing the observed number of annotated targets to the expected number based on the total number of genes in the genome. A one-tailed hypergeometric test was applied, and p-values were computed to assess statistical significance.

In addition, we separately evaluated enrichment in transcription factor targets that were up-regulated and down-regulated under drought conditions. For each TF, targets were partitioned according to the direction of their differential expression, and independent hypergeometric tests were performed for the up-regulated and down-regulated subsets. This allowed us to assess whether induced TFs preferentially regulate induced targets, and whether repressed TFs preferentially regulate repressed targets. The statistical framework was the same as described above, with one-tailed hypergeometric tests and p-value calculation for each subset. Results of this analysis are presented in Supplementary Figure 4.

### 2.7. Network Visualization

The merged gene regulatory networks were processed using custom R scripts and subsequently imported as text files into Cytoscape v3.10.3 for visualization (Shannon et al., 2003). To generate the TF interaction network, we first filtered the drought-responsive network to retain only interactions where both source and target nodes were TFs. From this TF-TF only subnetwork, we further filtered to include only the top-ranked TFs, as determined by their average scores across the three categories (upregulated, downregulated, and variable-response). This resulted in a subnetwork composed exclusively of predicted regulatory links between the top-ranked TFs. Node size was scaled according to outdegree (number of regulated TFs), and node color reflects the log2 fold-change under drought conditions. Visualization was refined using the Legend Panel tool to display node and edge attributes.

## 3. Results

### 3.1 An integrative transcriptomics approach identifies consistent drought-responsive genes

Identification of consistently regulated genes by drought is necessary because it enables the discovery of robust molecular responses that are maintained across different genotypes, tissues, and experimental conditions. Public repositories, such as the NCBI Sequence Read Archive (SRA), provide a valuable resource for accessing large-scale datasets from diverse experimental setups and bioprojects. We systematically searched the SRA database for RNA-seq libraries related to drought stress in *Vitis* species, retrieving datasets from multiple bioprojects. After rigorous curation, we retained only those with clear drought and control comparisons **(Figure 1).**

**Figure 1:**
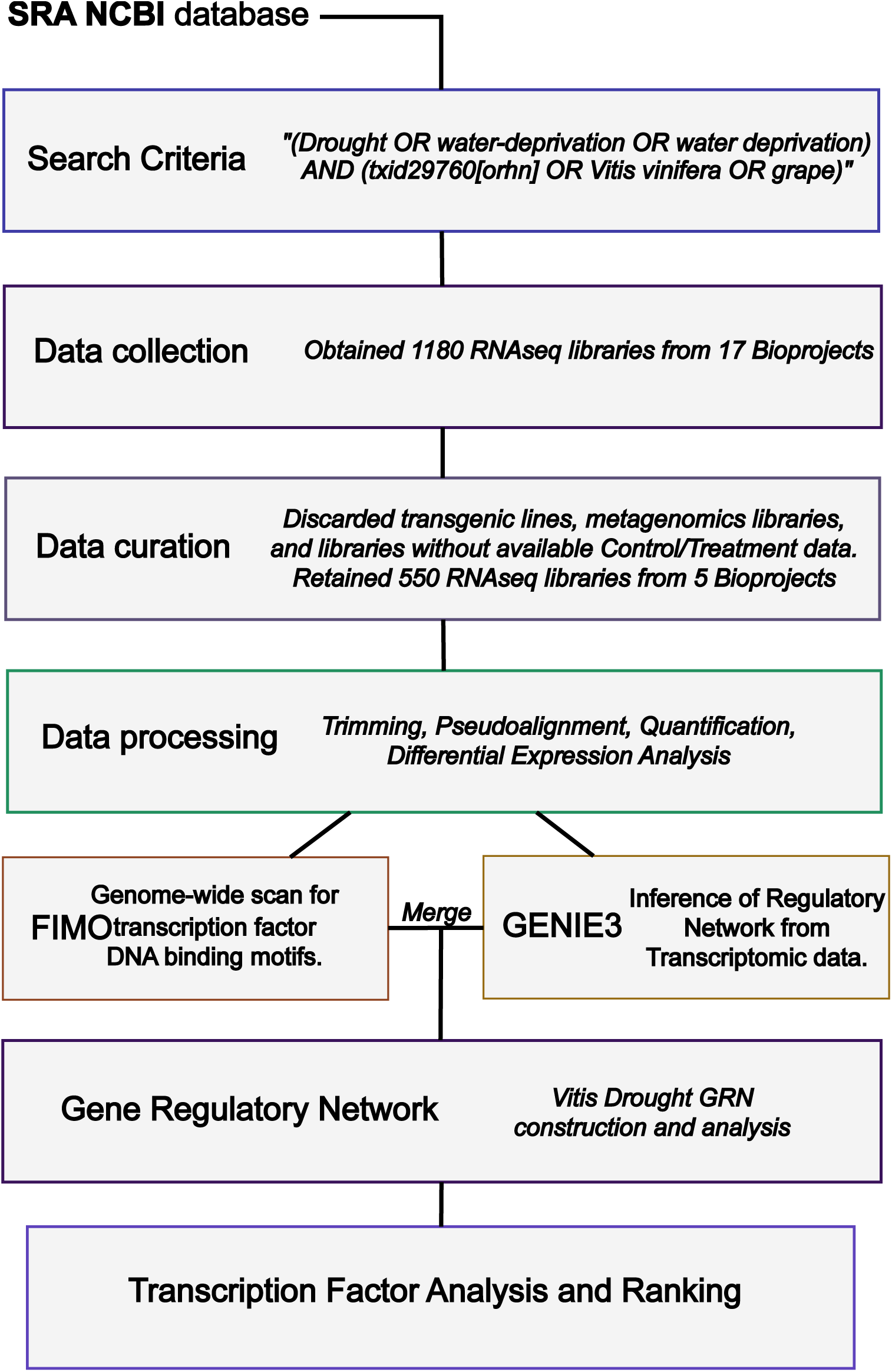
Workflow used to identify gene regulatory networks and key TFs responding to drought in V. vinifera and hybrid rootstocks. A total of 1180 RNA-seq libraries were retrieved from the NCBI SRA database. Libraries corresponding to transgenic lines, metagenomic samples, or lacking available information about control and treatment conditions were excluded, resulting in 550 libraries from 5 bioprojects. Data were processed through trimming, pseudoalignment, and quantification, followed by differential expression analysis. Genome-wide scanning for TF binding motifs and regulatory network inference were performed using FIMO and GENIE3. The resulting drought-responsive gene regulatory network was analyzed to prioritize TFs based on an average score that integrates magnitude of expression change, consistency across datasets, regulatory influence, and statistical validation. For variable-response TFs, the score was adjusted to account for their dynamic expression patterns, excluding magnitude of expression change.

Following data processing, differentially expressed genes (DEGs) were identified across experiments, capturing transcriptome responses to drought across different conditions, tissues, and genotypes. A total of 23 drought-control experiments from 5 bioprojects were included, and the datasets were named after the submitter’s last name or the first author of the associated publication (if available). These libraries represent a diverse range of genotypes, cultivars, and plant organs. The bioprojects and experiments, shown in **Figure 2**, include Zhu, Zheng, Khadka, Corso, and Botton. The results in **Figure 2.A** present all 23 transcriptomic comparisons grouped into five bioprojects, showing DEGs across various tissues, conditions, and time points. In the Zhu bioproject, conducted in leaves at 2, 4, and 8 days, up-regulated genes (red) consistently outnumber down-regulated genes (blue) across time points. Total number of DEGs increases over time. In the Zheng bioproject, also performed in leaves, fewer DEGs are observed compared to other bioprojects, with up-regulated genes nearly doubling the number of down-regulated genes. The Khadka bioproject evaluated roots and shoots at 14 days, where shoots show a higher number of DEGs, with down-regulated genes outnumbering up-regulated genes. In roots, up-regulated genes slightly predominate. The Corso bioproject included two genotypes, “M4” and “101-14” analyzed in leaves and roots at 2, 4, 7, and 10 days. In the “M4” rootstock genotype [(V. vinifera x V. berlandieri) x V. berlandieri cv Resseguier n.1] (Meggio et al., 2014), roots showed more up-regulated genes at 2 and 4 days. By 10 days, the number of DEGs dropped significantly, with up-regulated genes remaining more abundant. In leaves, up-regulated genes consistently exceeded down-regulated genes, and the total number of DEGs declined sharply by 10 days. In the “101-14” rootstock genotype (V. riparia x V. rupestris), roots exhibited a higher number of down-regulated genes across all conditions, while in leaves, this trend was observed only at 2 days under drought. Both roots and leaves experienced a reduction in total DEGs over time. Lastly, in the Botton bioproject, conducted in fruits at 9 days, down-regulated genes were more prevalent than up-regulated genes. The differential expression analysis across the five bioprojects revealed distinct patterns of up- and down-regulated genes under drought conditions, varying by tissue, genotype, and time point.

**Figure 2:**
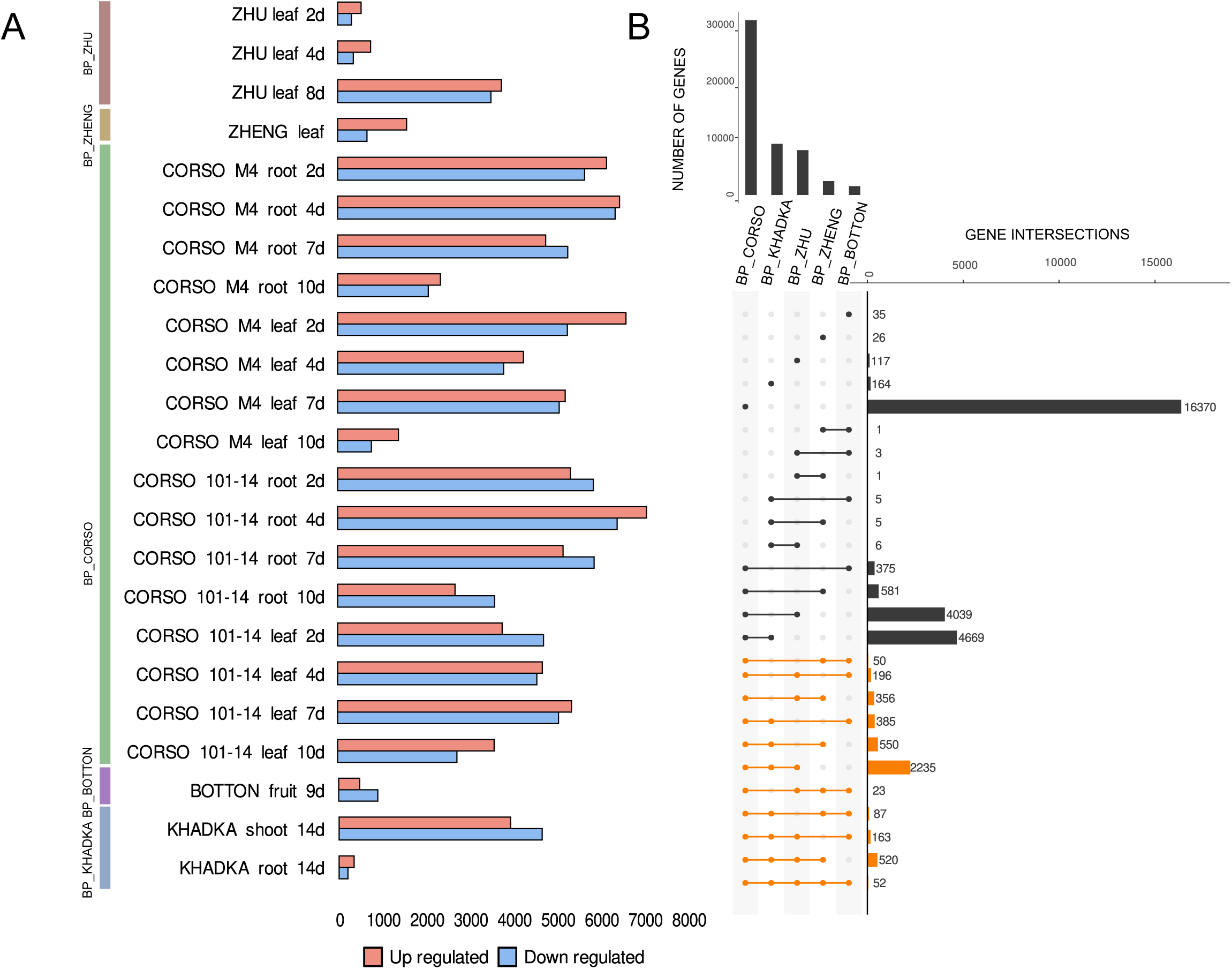
A core set of 4,617 drought-responsive genes emerges from the intersection of differentially expressed genes across multiple studies. **A)** Bar plots showing the number of upregulated (red) and downregulated (blue) genes across various bioprojects (BP) and experimental conditions, including different tissues (leaf, root, fruit, shoot) and time points under drought conditions. **B)** UpSet plot representing the intersections of differentially expressed genes among bioprojects. The bar plot on the top indicates the total number of differentially expressed genes per bioproject, while the horizontal bars on the right show the number of genes in each intersection. Orange lines highlight the genes that are part of the group enriched for the Gene Ontology (GO) term *Response to Water Deprivation*.

Despite these variations, a core set of drought-responsive genes was identified by analyzing the intersections of DEGs across bioprojects **(Figure 2.B)**. DEGs identified in at least three bioprojects were considered as consistent drought-responsive genes, resulting in a set of 4,617 genes (**Figure 3.A)**. The complete lists of DEGs identified for each individual comparison are available in Supplementary Table 2.

**Figure 3:**
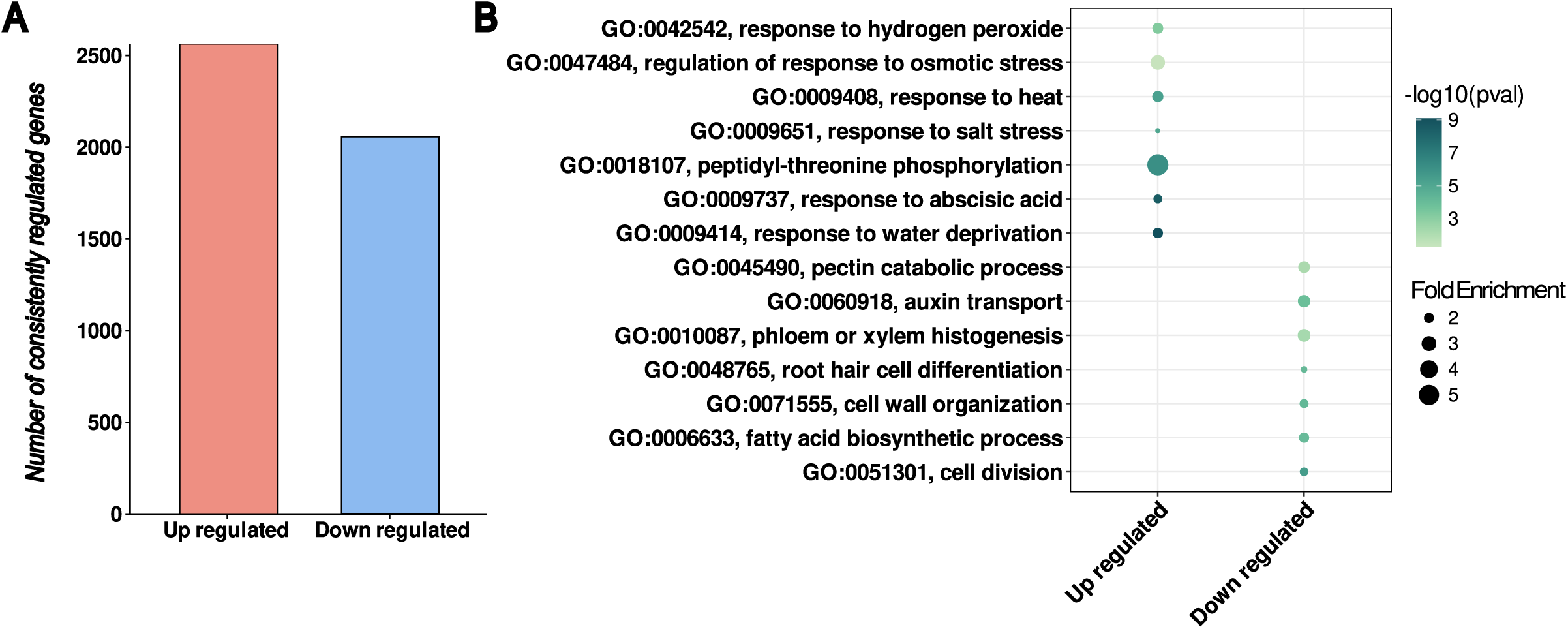
Upregulated genes under drought are predominantly associated with water deprivation and ABA responses, while downregulated genes are linked to growth-related processes. A) Bar plot showing the total number of consistently regulated genes for upregulated and downregulated categories. B) Gene Ontology term (Biological Process) enrichment analysis of Up and Down regulated genes under drought stress. The y-axis lists Gene Ontology (GO) terms associated with significantly represented biological processes. The x-axis distinguishes between up-regulated and down-regulated genes under the analyzed conditions. The size of the dots represents the degree of functional enrichment (FoldEnrichment), while the color indicates the level of statistical significance, measured as −log10(p-value) with darker tones denoting higher significance.

### 3.2 Consistently up regulated genes are strongly linked to water deprivation and ABA responsive pathways

Identifying the functional categories enriched among consistently regulated genes provides valuable insight into the molecular strategies grapevine employ to cope with water limitation. Of the 4,617 consistently regulated genes, 2,563 were up-regulated, while 2,054 were down-regulated under drought conditions **(Figure 3.A)**. Gene Ontology (GO) analysis of this gene set revealed distinct biological processes for both groups, with a significant overrepresentation of canonical drought-related biological processes among up-regulated genes (Yang et al., 2021). These processes included response to water deprivation, response to abscisic acid, response to salt stress, and response to heat stress, supporting the functional relevance of the identified drought-responsive genes. **(Figure 3.B; Supplementary Table 3).** In contrast, down-regulated genes were enriched in processes related to growth and cell division, such as phloem development, root hair cell differentiation, and cell wall organization or biogenesis (**Figure 3.B**).

These results underscore a clear functional dichotomy between the two groups: while up-regulated genes are predominantly involved in activating stress response mechanisms, down-regulated genes are associated with developmental and metabolic pathways typically repressed under drought. This sharp division reflects a coordinated transcriptional reprogramming in which distinct and discrete biological functions are either promoted or repressed.

### 3.3 Consistently drought-responsive genes reveal robust transcriptional signatures across organs and genotypes

After identifying consistent drought-responsive genes, analyzing their expression patterns across different experiments provides insights into shared regulatory mechanisms. **Figure 4** presents the expression profiles of the 4,617 consistent drought-responsive genes across 23 transcriptomic comparisons. The data is grouped by organ (leaf, root, fruit, shoot), genotype (*Vitis vinifera, Vitis riparia*, hybrids like 101-14, M4) and cultivar. Consistent transcriptome responses are evident, particularly in *Vitis riparia* shoots at 14 days, where both highly up- and down-regulated genes dominate the expression landscape. Time-course datasets, such as Corso and Zhu, reveal a progressive intensification of gene expression changes, peaking at 7 days in Corso and 8 days in Zhu, followed by a marked reduction at 10 days. Root tissues generally exhibit less pronounced responses compared to shoots, with some variability between genotypes i.e. 101-14 versus M4 and *V. riparia*. In leaves, consistent up-regulation trends are observed across genotypes, whereas responses in fruits and shoots highlight more diverse and condition-specific patterns. We prioritized genes that exhibit consistent regulation across multiple drought conditions and genotypes, emphasizing their robustness as drought-response markers. As an additional resource, we provide a supplementary dataset of strongly responsive genes, characterized by their high fold-change expression, which may serve as molecular markers for drought stress evaluation (Supplementary Figure 1, Supplementary Table 4). Furthermore, to facilitate a more detailed exploration of tissue-specific regulatory patterns, we present the expression profiles of genes uniquely regulated in each organ. This dataset offers insights into organ-specific drought responses, enabling a more targeted investigation of regulatory mechanisms in grapevine under drought conditions (Supplementary Figure 2; Supplementary Table 5). Their robustness under varying experimental conditions enhances their potential for further investigation as targets for improving drought tolerance in grapevine and hybrid rootstocks.

**Figure 4:**
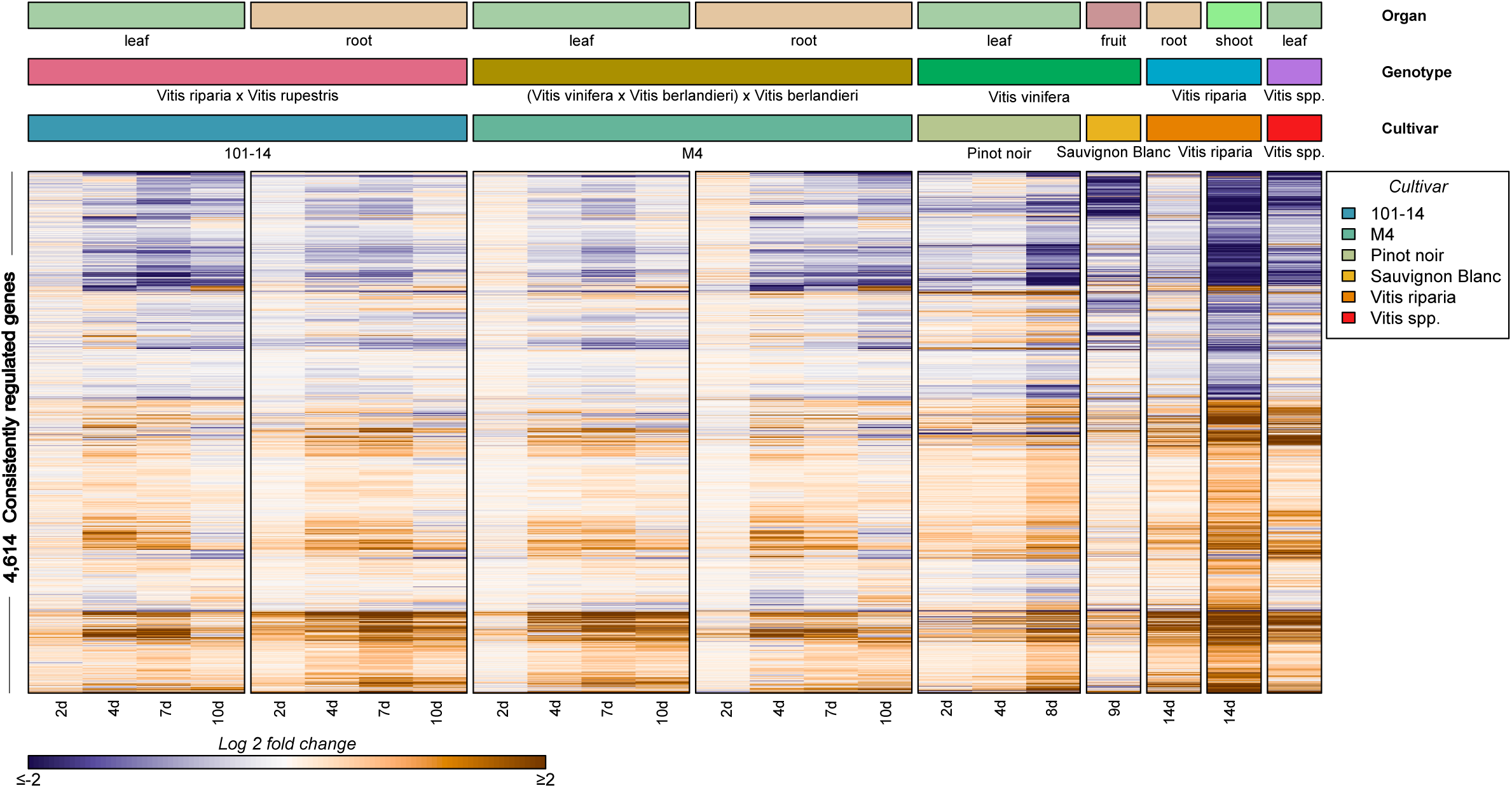
Consistently drought-responsive genes exhibit stable expression patterns across organs, genotypes, and experimental conditions. The heatmap displays the expression profiles of 4,617 genes consistently regulated under drought across 23 transcriptomic comparisons. Columns represent drought versus control contrasts, grouped by organ (leaf, root, fruit, shoot), genotype (*Vitis vinifera*, *V. riparia*, and hybrids such as 101-14 and M4), and cultivar. Rows correspond to individual genes. The color scale indicates log2 fold-changes (Log2FC), with red for upregulation and blue for downregulation.

### 3.4 Drought-responsive network comprise 5,058 nodes and 668 TFs

To better understand the regulatory mechanisms underlying drought response, we aimed to identify TFs with high regulatory potential within the drought-responsive gene network. First, we conducted a genome-wide motif analysis of promoter sequences using FIMO (Grant et al., 2011), a motif-based sequence analysis tool. We search for TF motifs for 1,388 TFs from 52 TF families obtained from the CIS-BP database (Catalogue of Inferred Sequences Binding Preferences, http://cisbp2.ccbr.utoronto.ca/) onto the *V. vinifera* promoter regions. The matches identified by FIMO predict regulatory links between TFs and their target genes, forming the basis of the GRN.

A second GRN was inferred using the GENIE3 machine learning algorithm, which identifies gene interactions based on expression data (Huynh-Thu et al., 2010). The final drought-responsive network was constructed by merging the FIMO-derived and GENIE3 networks (FIMO ⋂ GENIE3) and selecting all interactions involving the set of 4,617 consistent drought-responsive genes. This resulted in a drought-responsive network comprising 5,058 nodes and 178,960 interactions. The combination of FIMO-derived regulatory links with GENIE3’s network inference offers a robust framework for constructing reliable GRN (Mercatelli et al., 2020). The FIMO-only GRN, GENIE3-only GRN, merged GRN (FIMO ⋂ GENIE3), and the drought-responsive network basic statistics are shown in **Table 1**. Among the 5,080 nodes in the drought-responsive network, 668 are TFs. However, given the large number of candidates, it becomes essential to distinguish those with the highest regulatory influence.

**Table 1:** Gene regulatory networks statistics. The FIMO-only GRN, GENIE3-only GRN, merged GRN (Fimo ⋂ Genie3) and the drought-responsive network are described.

| Attribute | FIMO | GENIE3 | Merged Network | Drought-Responsive Network |
| --- | --- | --- | --- | --- |
| Nodes | 42,363 | 27,523 | 27,336 | 5,058 |
| Edges | 6,151,019 | 3,192,668 | 812,524 | 178,960 |
| TFs | 773 | 1155 | 670 | 668 |

### 3.5 Average score-based ranking reveals TFs with high regulatory potential

To identify the most relevant TFs, we applied an average scoring approach that integrates four criteria: (i) magnitude of transcriptomic response (log2FC), (ii) consistency across datasets, (iii) regulatory influence within the GRN (number of drought-responsive targets), and (iv) statistical enrichment for specificity of each TF. This strategy allowed us to prioritize TFs with the strongest regulatory potential in the drought response. An average score was calculated to integrate these four criteria to prioritize TFs with the highest regulatory influence.

Several of the top-ranked TFs identified through our average scoring strategy have been previously implicated in drought and ABA-mediated stress responses, supporting the validity of our approach. Notably, ABF2 (Nicolas et al., 2014) (also known as bZIP45), a key component of the ABA signaling pathway, was among the top-scoring TFs (J. Liu et al., 2014) (**Figure 5.A**). Other upregulated candidates included members of the Homeobox (HB) family, such as HB07, which is homologous to *AtHB7*, a well-established regulator of water stress (Y. Li et al., 2017). Additionally, MYB30A, member of the R2R3-MYB family, and NAC26, known to enhance drought tolerance in *A. thaliana*, were also highly ranked (Galbiati et al., 2011, Fang et al., 2016, Tello et al., 2015). The identification of these previously characterized TFs along with others like DREB2A (Aharoni et al., 2004; Khoudi, 2023) and SHN2 (Agarwal et al., 2006; Hou et al., 2023; Q. Liu et al., 1998), underscores the robustness of our approach and its ability to pinpoint both known and novel candidates for drought regulation.

**Figure 5:**
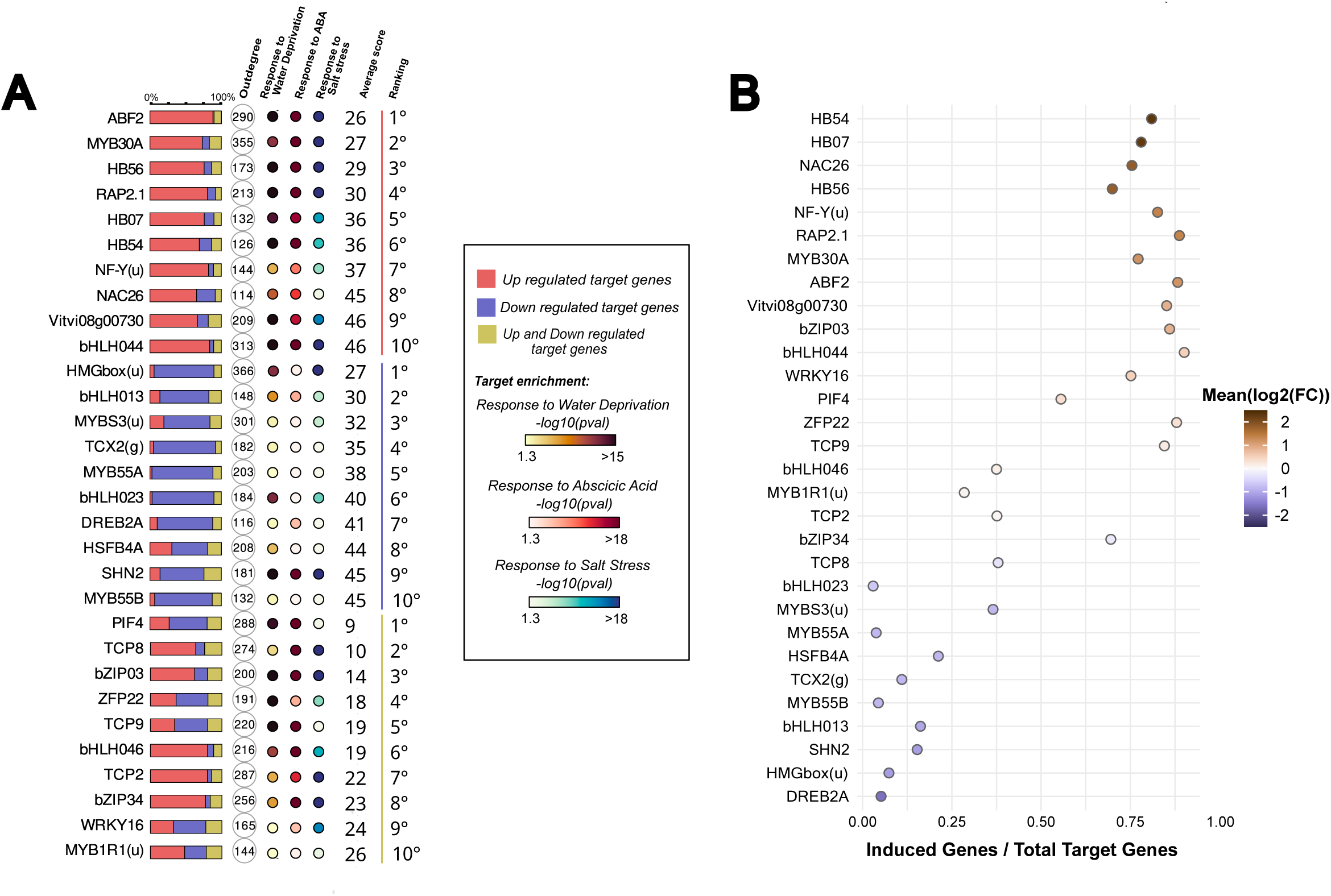
Top-ranked TFs display distinct regulatory profiles and are functionally linked to drought-related processes. A) Analysis of target genes regulated by top scoring TFs. Red bars indicate up-regulated proportion of targets, blue bars indicate down-regulated proportion of targets, and yellow bars indicate a proportion of genes with both up- and down-regulated expression. The total number of targets regulated by each transcription factor (outdegree) is displayed inside the circle next to the corresponding bar. Annotations indicating the enrichment of each TF’s target genes under Gene Ontology (GO) categories are also displayed; GO:0009414, Response to Water Deprivation and GO:0009737, Response to Abscisic Acid and GO:0009652, Response to Salt stress. Average score represents a composite measure used to rank each transcription factor, derived from the average rankings across multiple evaluation criteria. B) Distribution of the top scoring TFs in the drought-response network based on the ratio of induced to total target genes. Each point represents a TF, where the x-axis corresponds to the induced/total target gene ratio, and the y-axis lists TFs ranked by their transcriptomic response to drought. The color gradient reflects the log2 fold-change (Log2FC) of each TF, with red tones indicating upregulation and blue tones indicating downregulation.

**Figure 5.A** summarizes key regulatory features of the top 30 ranked TFs (10 per category: upregulated, downregulated, and variable). This figure integrates the total number of target genes per TF (outdegree), the proportion of targets that are upregulated, downregulated, or show variable responses, their GO enrichment in drought-related categories, and the average score used for ranking. Among the top TFs, several displayed high regulatory connectivity, including ABF2, MYB30A, bHLH044, HMGbox(u), and PIF4, each regulating a large number of drought-responsive genes. Notably, ABF2 regulated target genes enriched in all three GO categories, despite showing mild log2 fold-change values (Supplementary Figure 3). This highlights the strength of the average score in identifying regulators with significant downstream impact, even when their own transcriptomic response to drought is relatively modest.

Up-regulated TFs consistently showed increased expression across all experiments, with varying magnitudes of upregulation (Supplementary Figure 3). Target genes of such TFs were strongly enriched in GO categories related to water deprivation, abscisic acid, and salt stress responses. Downregulated TFs generally showed less enrichment in such GO categories compared to upregulated TFs. However, some exceptions include SHN2 and bHLH013, whose target genes were consistently enriched across all three stress-related GO terms. Additional TFs, such as bHLH23 and HMGbox(u), showed enrichment specifically in response to water deprivation and salt stress. Variable-response TFs showed mixed expression patterns, alternating between upregulation and downregulation depending on experimental conditions (Supplementary Figure 3). PIF4 and TCP9 were linked to genes highly enriched for water deprivation and abscisic acid responses, while bZIP03, bZIP34, bHLH046, and TCP2 consistently showed enrichment across all three stress responses.

### 3.6 TFs tend to establish regulatory interactions with others exhibiting the same expression direction in response to drought

A clear directional pattern was observed when examining the proportion of induced versus repressed targets. Induced TFs tended to activate a higher proportion of genes than they repressed, highlighting their potential role in activating drought-responsive genes (**Figure 5.A**). A similar pattern was observed among repressed TFs, where most of their target genes were downregulated. For instance, ABF2, an induced TF by drought, more than 75% of its targets were up-regulated, while a small proportion of targets were down-regulated.

To further explore the link between TF expression and regulatory output, we examined whether changes in TF expression levels correlate with the direction of regulation of their targets. As shown in **Figure 5.A**, TF expression patterns often mirrored the predominant response of their targets. **Figure 5.B** quantifies this relationship, plotting the proportion of induced targets relative to the magnitude of TF expression change. Strongly induced TFs, such as HB54, HB07, and NAC26, display ratios close to 1.0, indicating that nearly all of their targets are activated under drought stress. By contrast, TFs like DREB2A and SHN2 exhibit lower ratios, suggesting more limited or mixed regulatory effects. Overall, these results reveal a clear directional bias: induced TFs tend to activate their targets, whereas repressed TFs predominantly suppress them.

Supplementary Figure 4 presents the subset of TFs that are themselves differentially regulated under drought and examines the enrichment of their up- and downregulated targets. A clear pattern is observed: induced TFs tend to activate their targets, while repressed TFs predominantly suppress them. Notably, some TFs deviate from this trend by regulating targets in the opposite direction. These exceptions indicate that the observed directional concordance is not an artifact of the inference method, but rather a genuine feature of the drought-responsive regulatory network, where most TFs, but not all, act consistently with their own expression changes.

To assess whether the consistent expression trends observed between TFs and their target genes under drought stress could be explained by regulatory interactions among TFs themselves, we examined the structure and directionality of a TF-TF regulatory subnetwork. **Figure 6.A** shows the GRN of the top 30 drought-responsive TFs, comprising a total of 54 regulatory interactions. Central regulators with high outdegree values included MYB30A and ABF2, both upregulated under drought, while others such as SHN2 and HMGbox(u), though downregulated, also showed high connectivity. The subnetwork topology suggests a layered regulatory structure, with some TFs occupying hub-like positions.

**Figure 6:**
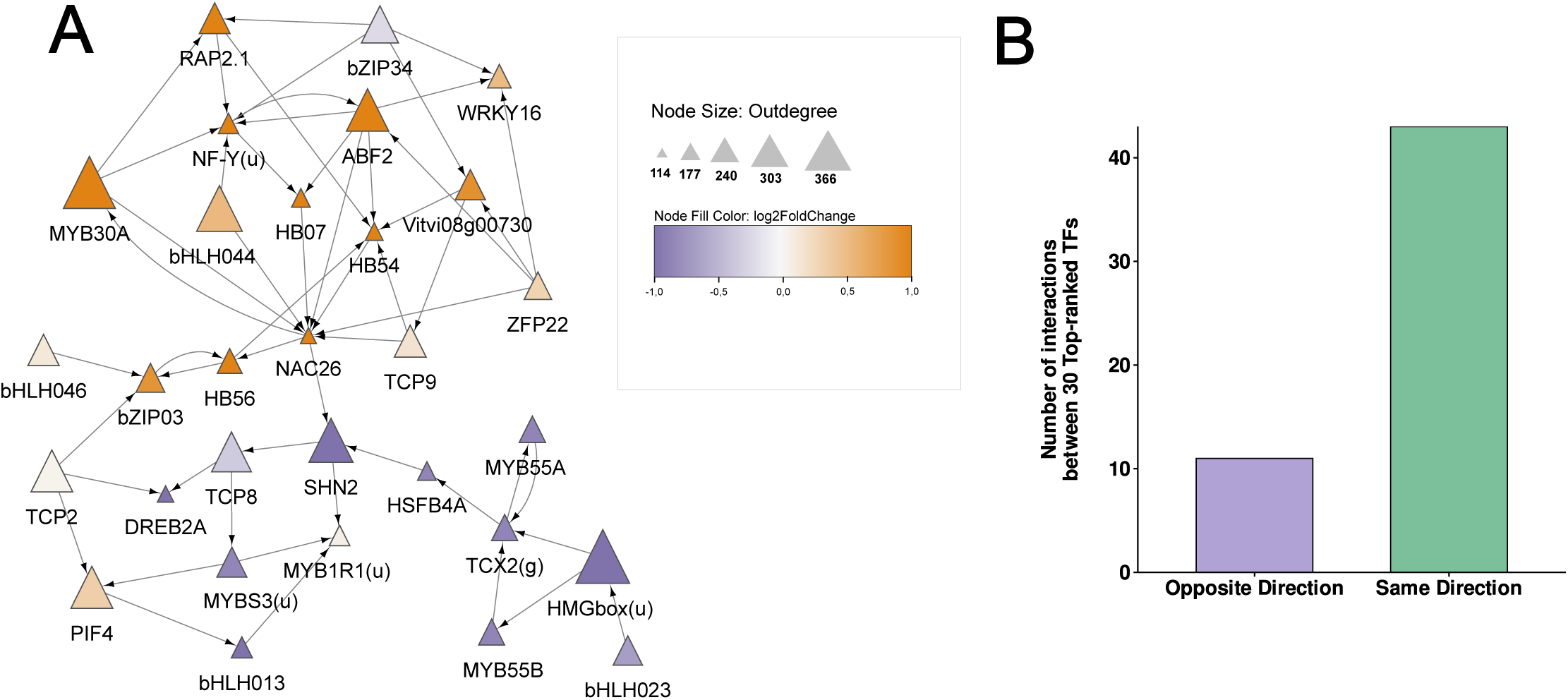
Regulatory interactions between TFs are predominantly established between TFs with the same direction of expression change under drought stress. **A)** Regulatory network comprising the top 30 TFs responsive to drought conditions. Each node represents a TF, with node size proportional to its outdegree (i.e., the number of downstream TFs it regulates). Node color indicates the log2 fold-change (Log2FC) in expression under drought, with red shades representing upregulation and purple shades representing downregulation. Directed edges indicate predicted regulatory interactions between TFs. **B)** Bar plot quantifying the number of regulatory interactions classified by expression directionality. “Same direction” interactions refer to those where both TFs are either upregulated or downregulated. “Opposite direction” interactions occur when one TF is upregulated and the other is downregulated, regardless of source or target orientation.

To investigate whether TFs tend to regulate other TFs with similar expression responses, we classified the 54 interactions based on the relative direction of expression change between connected TFs. **Figure 6.B** shows that the majority of regulatory interactions occurred between TFs that were either both upregulated or both downregulated under drought conditions. Only a minority of interactions involved TFs with opposite expression trends. These results support the idea that the observed consistency in expression between TFs and their downstream targets may be partially explained by hierarchical regulation among similarly responsive TFs, where drought-activated TFs regulate downstream TFs that are also upregulated.

## 4 Discussion

In this study, we performed a comprehensive transcriptomic meta-analysis of *V. vinifera* and hybrid rootstocks under drought conditions across multiple organs (leaves, roots, berries, and shoots) and genotypes, by integrating data from 550 libraries derived from five independent bioprojects. This allows us the inspection of consistently regulated genes, as a resource, and a data-rich opportunity to predict gene regulatory networks and identify relevant candidate TFs to modulate drought responses in *V. vinifera*.

The integration of a large and heterogeneous transcriptomic dataset is an advance, as provides a framework to identify genes consistently regulated by drought stress across diverse biological contexts. This approach enables the discovery of high-confidence drought-responsive genes while accounting for variation in experimental designs, genotypes, and tissues. Similar strategies have been successfully applied in *Arabidopsis thaliana*, where meta-analyses have shown that a significant proportion of DEGs overlap with previously characterized stress-responsive genes (Sharma et al., 2018), reinforcing the utility of this methodology.

In our study, we curated 550 RNA-seq libraries from five independent bioprojects, encompassing 23 drought vs. control comparisons across leaves, roots, berries, and shoots in multiple genotypes. To address potential technical bias between studies, we aggregated comparisons within each bioproject. For those including time-course experiments, DEGs from all-time points were collapsed into a single comparison, with the aim of reducing redundancy and improving consistency. Despite the diversity in tissues and experimental conditions, we identified a core set of 4,617 genes consistently responsive to drought, defined as DEGs present in at least three different bioprojects. This threshold was established through enrichment analysis of related biological GO terms (see Methods 2.2 and Supplementary Table 1) aiming for a balance between stringency and inclusiveness.

A large fraction of the 4,617 drought-responsive genes showed similar expression patterns across distinct tissues and genotypes, indicating a conserved transcriptional response to drought in *V. vinifera*. Similar convergence has been reported in other species. In *Arabidopsis thaliana*, over 2,500 and 3,600 DEGs have been reported in roots and shoots, respectively, with several hundred genes showing co-regulation across tissues (Ghorbani et al., 2019; Shariatipour & Heidari, 2018). Cross-species meta-analyses have also identified conserved components: 225 drought-regulated genes shared among *Arabidopsis*, rice, wheat, and barley (Shaar-Moshe et al., 2015), and 69 conserved genes across tolerant and sensitive cereal genotypes (Baldoni et al., 2021). Additional studies in wild soybean, sugarcane, and pecan report hundreds to thousands of DEGs, with subsets showing consistent expression between genotypes (Aleem et al., 2021; P. Li et al., 2022; Zhu et al., 2023). These findings indicate that drought triggers a core transcriptional response that is at least partially conserved across tissues, genotypes, and species. The consistent patterns observed in *Vitis* support the robustness of the identified gene set and its potential value for comparative studies and as a source of candidate markers or regulators for improving drought tolerance.

GRN inference is a tool for understanding transcriptional drought responses in *V. vinifera*, particularly given the limited availability of experimentally validated TF-target interactions compared to model species like *Arabidopsis thaliana* (Brooks et al., 2021). The drought-responsive GRN inferred in this study comprises 5,058 nodes and 178,960 regulatory interactions, including 668 TFs. This represents a larger set of TFs and a more focused network than those reported in *Arabidopsis thaliana* under similar conditions, where studies have typically identified between 117 and 261 TFs (Akhtartavan & Talebi, 2018; Kumar et al., 2023). While the Arabidopsis Drought Gene Co-expression Network (ADGCN) described by Kumar et al. (2023) includes over 400,000 average edges derived from gene correlation, our network presents fewer interactions. This difference likely reflects methodological contrasts: our integrative approach combines GENIE3 inference with motif-based evidence from FIMO, which may reduce the inclusion of indirect or spurious co-expression links by restricting predictions to TFs with putative binding sites in their targets.

The relationship between TF expression levels and the proportion of induced target genes suggests that induced TFs play a role in activating downstream regulatory networks in response to drought stress. It is plausible that these TFs, upon induction, activate not only their direct targets but also other positive regulators involved in the stress response. This could explain the higher proportion of induced target genes observed for these TFs, as they likely promote the activation of key protective and adaptive mechanisms such as osmotic regulation, cellular protection, and stress signaling. GO enrichment analysis of the genes consistently induced under drought supports this interpretation, with terms such as *response to water deprivation* and *abscisic acid signaling* among the most significantly overrepresented, reflecting a broad reprogramming of gene expression toward stress adaptation.

It is important to note that GENIE3 infers regulatory relationships by modeling gene expression with ensembles of regression trees, ranking TFs according to their predictive contribution for each target gene. One might argue that the concordance we observe between TF expression and the direction of target gene regulation could simply result from co-expression patterns captured by this approach. However, GENIE3 does not intrinsically differentiate between transcriptional activation and repression. Independent functional assays in *Arabidopsis* provide evidence that this concordance reflects genuine regulatory interactions. ABI5, an induced TF by drought, has been validated by mutant analyses as a positive regulator that activates its downstream targets (Kumar et al., 2023). Overexpression of the ABA-responsive TF ABF3 induces canonical drought and ABA-inducible target genes (Abdeen et al., 2010). Complementing this, RNA-seq assays using the cell-based TARGET system, which enables the identification of genes directly induced or repressed by specific TFs, confirmed that several ABA-pathway TFs, including ABF1, ABF3, and HB7, predominantly activate their downstream targets. These direct targets are significantly enriched in ABA-responsive genes (Brooks et al., 2021), reinforcing the notion that drought-activated TFs largely function as inducers within stress-responsive networks.

Consistently, in grapevine, overexpression of VvABF2 followed by transcriptome profiling demonstrated that the majority of its regulated genes were upregulated under ABA treatment treatment (Nicolas et al., 2014) providing further evidence that ABA-related TFs primarily act as transcriptional activators during drought responses.

Conversely, RNA-seq assays using the TARGET system also identified TFs with predominantly repressive activity. For example, ERF056, HSFB2A, and ZFP4 were found to repress the majority of their downstream targets (Brooks et al., 2021). Consistent with this pattern, independent studies have demonstrated that ZFP8, a close member of the same family of C2H2 zinc-finger TF, is downregulated by ABA during drought responses (Tian et al., 2024). These observations support that the directional trends identified here are not artifacts of the inference method but reflect underlying biological regulation.

Nevertheless, the prevalence of directional concordance raises the question of how negative regulators fit into drought-responsive networks. According to our criteria, such TFs should be induced while repressing a large proportion of their targets. Target enrichment analysis confirmed that such cases exist, but they are much less abundant compared to TFs that regulate targets in the same direction. A set of TFs with this pattern was identified in the network, although they did not rank among the top regulators based on average score. These TFs show less consistent expression patterns across the full dataset and do not exhibit strong expression changes, raising the hypothesis that their identification and interpretation may require tissue-specific analyses rather than broad, diverse datasets. Exploring their regulatory mechanisms could be the focus of future studies in grapevine, contributing to a deeper understanding of the diversity of regulatory responses to drought.

The TF regulatory network inferred from the top 30 drought-responsive TFs reveals that most regulatory interactions occur between TFs with similar expression trends under drought stress. In this network, TFs such as MYB30A and ABF2, both upregulated, show high outdegree values, indicating a prominent role in regulating other TFs. Conversely, SHN2 and HMGbox(u), despite being repressed under drought, also display extensive connectivity, suggesting that transcriptional repression does not preclude regulatory influence.

To evaluate whether TFs tend to regulate others with similar expression, we classified the 54 predicted interactions based on the expression direction of the source and target TFs. Most interactions occurred between TFs that were either both upregulated or both downregulated, while interactions between TFs with opposing expression trends were comparatively rare.

This pattern supports the idea that drought-responsive TFs are preferentially connected to others exhibiting the same direction of regulation. Induced TFs tend to activate other induced TFs, potentially reinforcing stress-related transcriptional programs. Similarly, repressed TFs are more likely to be connected to other downregulated regulators, possibly coordinating the suppression of non-essential pathways such as growth and energy-intensive metabolism (Haghpanah et al., 2024;Nour et al., 2024). The relative scarcity of opposite-direction interactions suggests that transcriptional responses under drought are structured to favor same-direction regulatory interactions. Additionally, the presence of TFs with markedly different levels of connectivity highlights a hierarchical organization, in which some TFs act as regulatory hubs while others occupy more limited or context-specific roles, as previously demonstrated for ABA responses (Song et al., 2016). This structured architecture likely contributes to the stability and specificity of the transcriptional response under water stress.

Several of the TFs prioritized in our drought-responsive network belong to families previously implicated in abiotic stress regulation, including AP2/ERF, bZIP, MYB, and bHLH. This is consistent with findings in *A. thaliana*, where these TF families were also reported as abundant under drought and cold stress conditions (Sharma et al., 2018). These families are known to participate in hormone signaling, osmotic stress response, and growth regulation, supporting the functional relevance of our prioritized TFs.

The hierarchical organization observed in our network, with certain TFs acting as highly connected hubs, provides a strategic framework for prioritizing candidates for experimental validation. Notably, some of these TFs, such as ABF2, NAC26, and HB07, have been previously linked to drought tolerance in other species (Fang et al., 2016; J. Liu et al., 2019; Olsson et al., 2004), reinforcing their potential roles in grapevine. ABF2 is a well-characterized master regulator of ABA-dependent transcriptional responses and has been shown to directly activate downstream genes and other TFs involved in drought adaptation (Song et al., 2016; Yoshida et al., 2014). NAC26 and HB07 have also been associated with stress-induced reprogramming and hormonal signaling, and both have been identified as central nodes in drought-responsive GRNs, supporting their potential roles as master TFs in the regulatory hierarchy (Fang et al., 2016; Olsson et al., 2004; Ré et al., 2014).

Given that transcriptional regulation often depends on the combined action of multiple TFs, integrating complementary approaches such as chromatin accessibility profiling (ATAC-seq) and TF-binding assays (ChIP-seq or DAP-seq) will be important to identify direct regulatory interactions (Song et al., 2016). Functional validation through overexpression or genome editing (e.g., CRISPR-Cas9) will help confirm their regulatory roles and assess their utility for improving drought resilience.

However, translating these findings into applied strategies requires consideration of physiological trade-offs. Enhancing drought tolerance without compromising yield remains a major challenge. In this context, targeting TFs that modulate water-use efficiency or coordinate ABA signaling and growth restraint may offer promising avenues (Tardieu, 2022; Vadez et al., 2024).

## 5 Conclusion

This study provides a comprehensive analysis of drought-responsive gene regulation in *V. vinifera* and hybrid rootstocks. By integrating transcriptomic data from multiple bioprojects, we identified a core set of 4,617 genes consistently regulated under drought conditions. Using integrative genomics approaches, including promoter motif analysis and machine learning-based network inference, we constructed a high-confidence drought-responsive gene regulatory network and identified key TFs, such as ABF2 and MYB30A, as central regulators.

A major finding is the directional regulatory influence within the network: upregulated TFs predominantly activate upregulated target genes, while downregulated TFs are mainly associated with repression of downregulated targets. This suggests a coordinated transcriptional control strategy that amplifies specific responses to drought, either activating protective mechanisms or repressing growth-related processes.

These insights provide valuable targets for future functional validation and genetic improvement efforts aimed at enhancing drought tolerance in grapevine. Further experimental work will be essential to confirm the regulatory roles of these TFs and their potential applications in crop improvement.

## 6 Supplementary Data: Tables and figures

**ST1: Enrichment analysis of water deprivation-related DEGs across grouped bioprojects.** The table presents the results of a hypergeometric test applied to three different bioproject thresholds (≥3, ≥4, and ≥5) to identify which group is most enriched with differentially expressed genes (DEGs) annotated under the GO term GO:0009414, Response to water deprivation. The table shows the total number of DEGs in each group, the number of DEGs annotated with the water deprivation GO term, and the corresponding p-values for each threshold.

**ST2: Differentially expressed genes per comparison.** This file contains the complete output of DESeq2 analyses. Each of the 23 worksheets corresponds to one comparison and includes gene identifiers, log2 fold change, statistical significance values, and other metrics generated by DESeq2.

**ST3. Gene Ontology (GO) enrichment analysis of 4,617 consistent drought-responsive genes.** This file contains the results of GO term enrichment performed with topGO: summary of significantly enriched GO terms across conditions; complete enrichment results for up-regulated genes; complete enrichment results for down-regulated genes; and the list of the 4,617 consistent drought-responsive genes used in the enrichment analysis.

**ST4. Highest fold change drought-responsive genes.** This file contains genes with the strongest transcriptional responses to drought. It includes (i) the mean log2 fold change across all experiments, showing the most strongly up- and down-regulated genes, and (ii) the complete fold change matrix from 23 comparisons, with per-gene values across experiments and the number of experiments in which each gene was differentially expressed.

**ST5. Tissue-specific drought-responsive genes.** This file contains the genes identified as root-specific (Root) and leaf-specific (Leaf) according to their expression patterns under drought conditions.

**SF1: Heatmap of highest fold change drought-responsive genes.** The heatmap displays the top 10 drought-responsive genes with the highest log2 fold changes in expression. Rows represent individual genes, while columns correspond to specific libraries and the “Average Expression” column, which summarizes the mean log2 fold change across all libraries and is distinguished from individual library data to provide an overview of the overall gene response. This visualization highlights the differential expression patterns of the most drought-responsive genes across libraries.

**SF2: Heatmap of tissue-specific genes: Root vs Leaf.** Heatmaps display the expression profiles (log2 fold change, drought vs. control) of genes classified as tissue-specific. Left: leaf libraries; Right: root libraries. Rows represent root-specific or leaf-specific genes, while columns correspond to libraries grouped by bioproject. Color scale indicates the level of induction (orange) or repression (blue). This visualization highlights differential expression patterns between root- and leaf-specific genes under drought conditions.

**SF3: Top ranked Transcription Factors regulating the drought-responsive network.** Heatmap displaying the expression profiles of top scoring transcription factors across all experiments. Columns were grouped by tissue and rootstocks/cultivars, and TFs were grouped by ranking (Variable-response TF, Up-regulated TF, and Down-regulated TF). The heatmap includes annotations indicating the enrichment of each TF’s TARGET genes under Gene Ontology (GO) categories; GO:0009414, Response to Water Deprivation and GO:0009737, Response to Abscisic Acid and GO:0009652, Response to Salt stress. The scale bar reflects Log 2 fold changes, with red indicating upregulation and blue indicating downregulation. Gene symbols were primarily obtained from the Gene Reference Catalogue (Vitiviz); genes marked with (u) were retrieved from UniProt, and those marked with (g) were named by Gramene.

**SF4: Directionality of target enrichment relative to transcription factor expression under drought conditions.** A) Enrichment of up-regulated target genes (y-axis, log2 scale) as a function of the transcription factor log2 fold change (x-axis). B) Enrichment of down-regulated target genes relative to TF log2 fold change. Each point corresponds to a TF, colored by the statistical significance of the enrichment (-log10(padj)); warmer colors indicate higher significance. Density contours show the distribution of TFs, and marginal plots display the frequency of TF log2 fold changes. Dashed horizontal lines represent no enrichment, and dashed vertical lines represent no change in TF expression. Most TFs are located in quadrants reflecting concordance between their regulation and the regulation of their targets, while fewer TFs appear in the opposite quadrants, showing cases of discordant regulation.

