## Supplementary figures and images for "Consistent drought regulation in grapevine is driven by directional transcription factor activity"

### Supplementary Figure 1

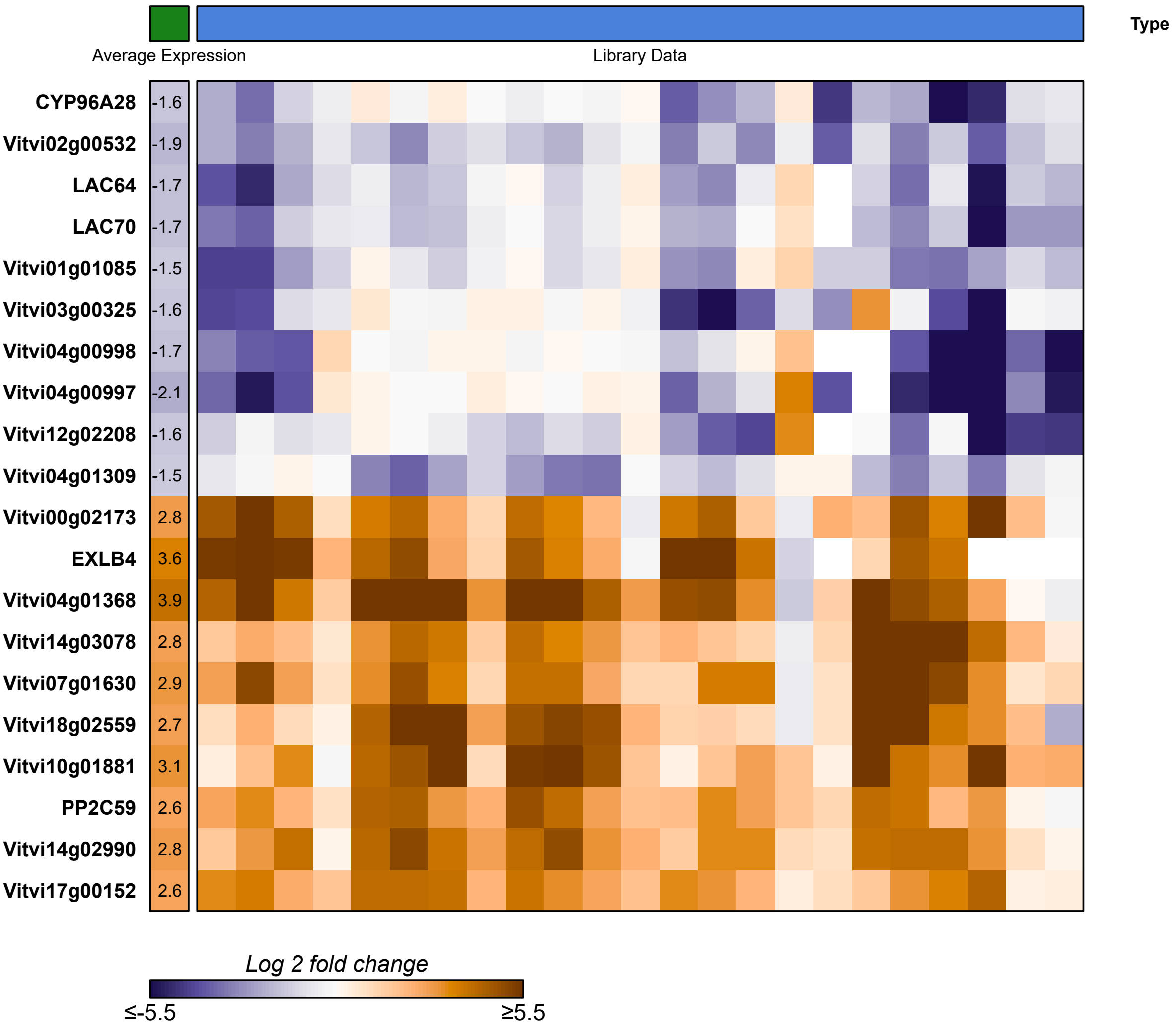

### Supplementary Figure 2

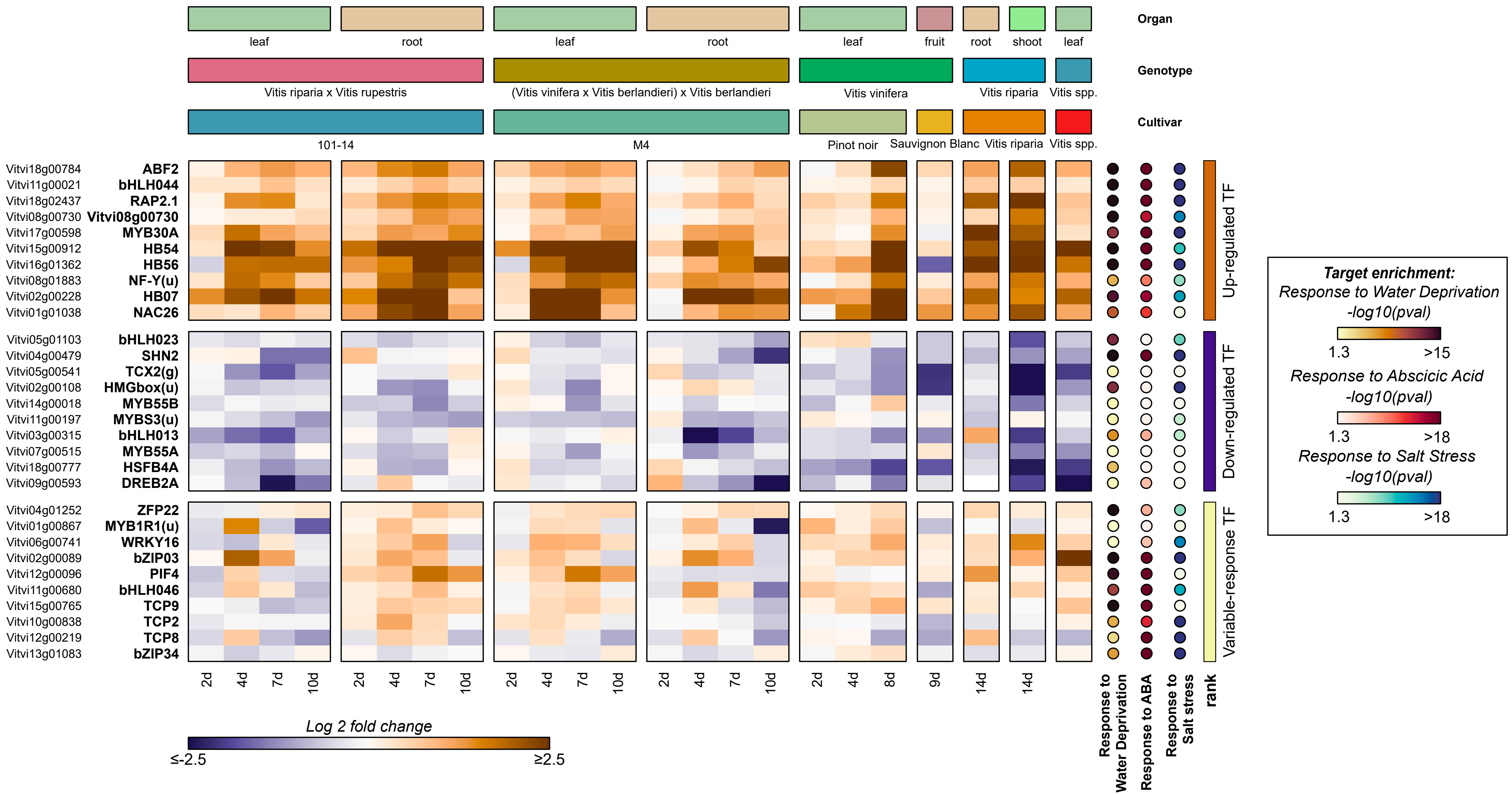

### Supplementary Figure 3

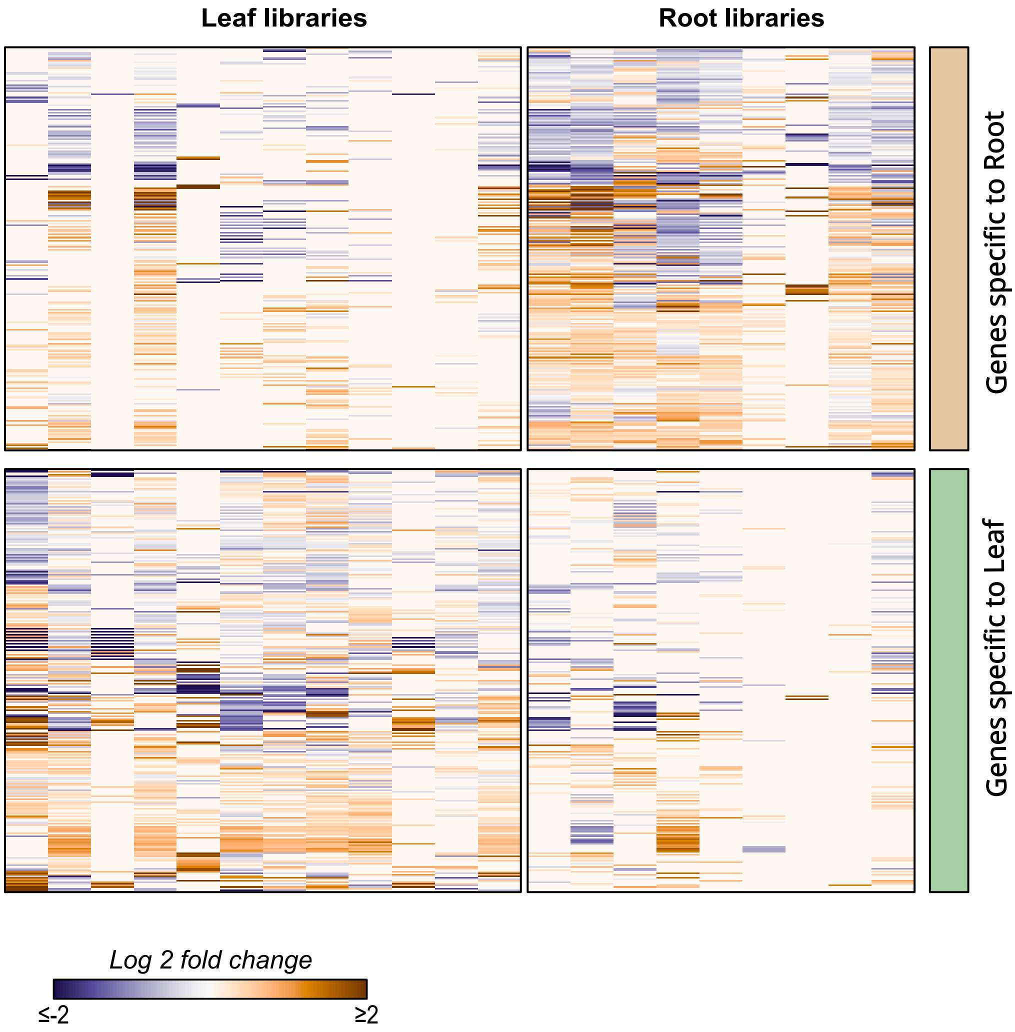

### Supplementary Figure 4

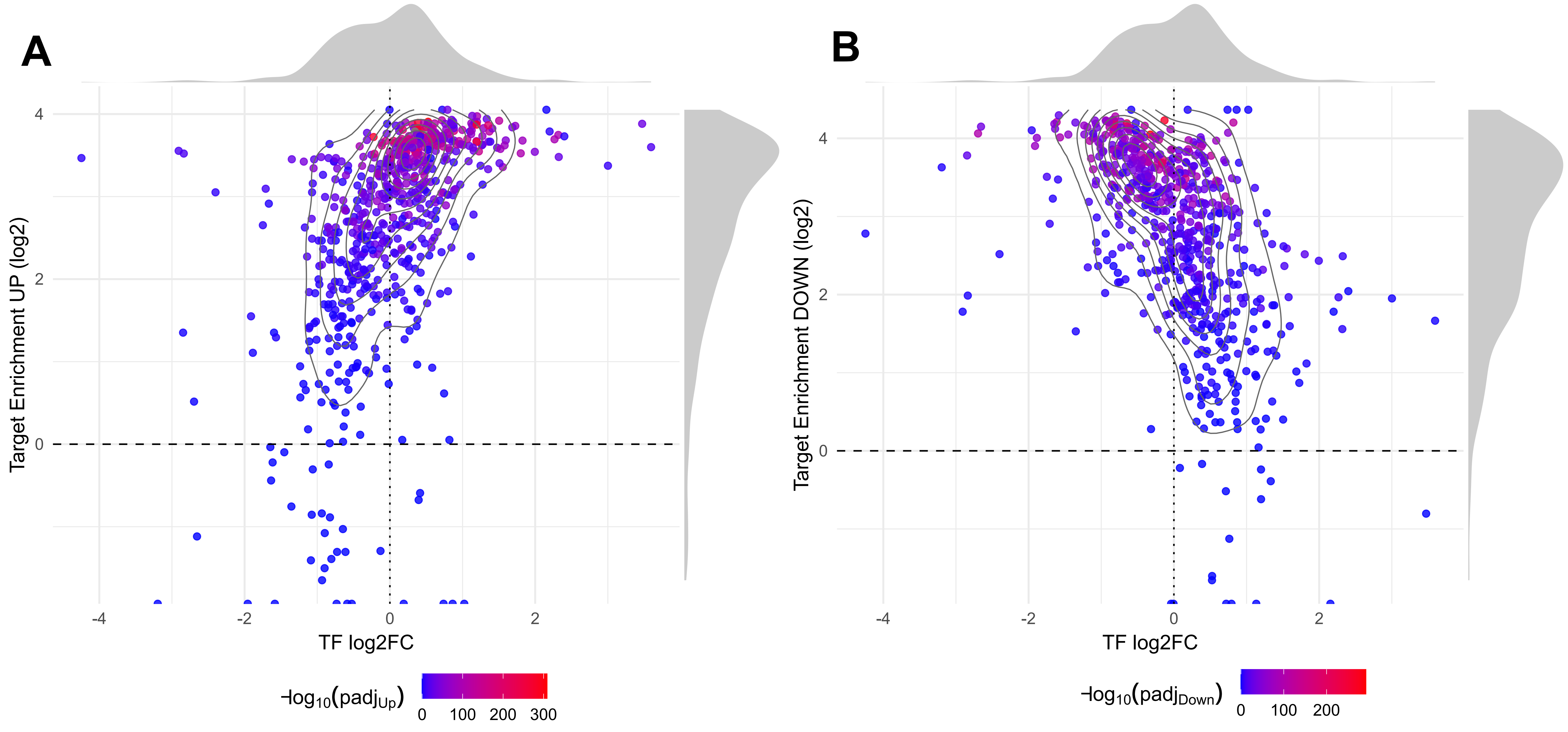
